# Characterization of β-lactamase and virulence genes in *Pseudomonas aeruginosa* isolated from clinical, environmental and poultry sources in Bangladesh

**DOI:** 10.1101/2023.12.16.572010

**Authors:** Raihana Islam, Farhana Binte Ferdous, M. Nazmul Hoque, Nowshad Atique Asif, Md. Liton Rana, Mahbubul Pratik Siddique, Md. Tanvir Rahman

## Abstract

The emergence and spread of multidrug-resistant pathogens like *Pseudomonas aeruginosa* are major concerns for public health worldwide. This study aimed to investigate the prevalence of circulating *P. aeruginosa* isolated from clinical, environmental and poultry sources in Bangladesh, their antibiotic susceptibility, β-lactamase and virulence gene profiling using standard molecular and microbiology techniques. We collected 110 samples from five different locations, *viz*., BAU residential area (BAURA; n=15), BAU Healthcare Center (BAUHCC; n = 20), BAU Veterinary Teaching Hospital (BAUVTH; n=22), Poultry Market (PM; n=30) and Mymensingh Medical College Hospital (MCCH; n=23). After overnight enrichment in nutrient broth, 89 probable *Pseudomonas* isolates (80.90%) were screened through selective culture, gram-staining and biochemical tests. Using genus- and species-specific PCR, we confirmed 22 isolates (20.0%) as *P. aeruginosa* from these samples. Antibiogram profiling revealed that 100.0% *P. aeruginosa* isolates (n = 22) were multidrug-resistant isolates, showing resistance against Doripenem, Penicillin, Ceftazidime, Cefepime, and Imipenem. Furthermore, resistance to aztreonam was observed in 95.45% isolates. However, *P. aeruginosa* isolates showed a varying degree of sensitivity against Amikacin, Gentamicin, and Ciprofloxacin. The *blaTEM* gene was detected in 86.0% isolates, while *blaCMY*, *blaSHV* and *blaOXA,* were detected in 27.0%, 18.0% and 5.0% of the *P. aeruginosa* isolates, respectively. The *algD* gene was detected in 32.0% isolates, whereas *lasB* and *exoA* genes were identified in 9.0% and 5.0% *P. aeruginosa* isolates. However, none of the *P. aeruginosa* isolates harbored *exoS* gene. Thus, this study provides novel and important data on the resistance and virulence of *P. aeruginosa* currently circulating in clinical, environmental and poultry environment of Bangladesh. These data provide important insights into the emergence of β-lactamase resistance in *P. aeruginosa*, highlighting its usefulness in the treatment and control of *P. aeruginosa* infections in both humans and animals.

## 1. Introduction

*Pseudomonas aeruginosa*, a Gram-negative bacterium, has gained prominence as an emerging opportunistic pathogen with major clinical consequences, particularly in hospital settings where it poses a large risk to immunocompromised persons [1, 2]. *P. aeruginosa*, known for its adaptability and metabolic diversity, is a prominent cause of nosocomial infections, producing severe acute and chronic illnesses in individuals with a variety of vulnerabilities [3, 4]. Its ubiquity goes beyond healthcare facilities since it thrives in a variety of habitats, including soil, water, and even oil-polluted environments, giving it the term "ubiquitous" or "common soil and water bacterium" [5].

Antimicrobial resistance (AMR) in *P. aeruginosa*, is a significant concern in healthcare settings due to its ability to cause severe infections, particularly in individuals with weakened immune systems. *P. aeruginosa* is inherently resistant to many antibiotics and has a remarkable capability to develop further resistance mechanisms [6]. The increasing fear of *P. aeruginosa* is exacerbated by the worldwide problem of antimicrobial resistance (AMR), which has been dubbed a "crisis of the twenty-first century" [7]. Antibiotic overuse has developed multidrug resistance among bacteria, such as *P. aeruginosa*, posing a severe danger to public health and economies, particularly in low and middle-income countries (LMICs) in Africa and Asia, including Bangladesh. The growth in antibiotic-resistant *P. aeruginosa* strains, with over 10% of worldwide isolates demonstrating multidrug resistance, adds to the difficulties in treating infections caused by this bacterium [8]. Notably, *P. aeruginosa* uses a variety of resistance mechanisms, including gene expression under stress and the production of antibiotic-resistant biofilms, which reduces the efficiency of traditional therapies [9]. Antibiotic-resistant *P. aeruginosa* is found in various habitats, including wastewater and soils, emphasizing the necessity of studying the frequency and resistance patterns of these clinically relevant bacteria in various contexts [10]. The recent development of extended-spectrum beta-lactamases (ESBLs) complicates the clinical situation even more, with ESBLs being a major contributor to bacterial resistance worldwide [11]. As a result of the complications imposed by ESBL-producing isolates, microbiologists, medics, and scientists working on creating novel antimicrobial medications face extra hurdles [11, 12].

*P. aeruginosa* is known for its diverse array of virulence factors, contributing significantly to its pathogenicity and ability to cause infections. A variety of virulence factors have been discovered in *P. aeruginosa* that can significantly contribute to the pathogenicity of this bacterium [4, 13]. Toxin A (*toxA*), exotoxin A (*exoA*), alkaline protease (*aprA*), elastase, and exoenzymes (S, U, and T *exoS*, *exoU*, *exoT*) are the principal virulence factors. *LasB*, a zinc metalloprotease, exhibits elastolytic activity specifically on lung tissue of human [14]. Chronic lung infections are significantly influenced by the *algD* gene, which is accountable for the alginate capsule of *P. aeruginosa* [15]. In addition, exotoxin A is secreted via Type II secretion systems, thereby contributing to the extracellular pathogenicity of the bacterium [16]. The presence and interplay of these virulence factors in *P. aeruginosa* contribute to its pathogenicity, enabling it to cause a wide range of infections, particularly in immunocompromised individuals or those with underlying health conditions. Understanding these virulence mechanisms is crucial for developing targeted strategies to combat *P. aeruginosa* infections and improve patient outcomes [4].

While most *P. aeruginosa* research in Bangladesh has concentrated on antibiotic resistance profiles in clinical and wastewater isolates, but a thorough examination of environmental isolates in the specified region is lacking. Thus, this study aimed to provide significant insights into the prevalence, antimicrobial resistance, and virulence profiles of multidrug-resistant *P. aeruginosa* isolates from clinical, environmental and poultry sources in some selected areas of Bangladesh. The findings of this study can significantly contribute to our knowledge of the dynamics of *P. aeruginosa*, which in turn can inform public health policies, clinical practices, and future research efforts aimed at controlling infections caused by this pathogen.

## 2. Materials and Methods

### 2.1 Isolation and identification of *P. aeruginosa*

This cross-sectional study was conducted in the Bacteriology Laboratory of the Department of Microbiology and Hygiene, Bangladesh Agricultural University (BAU), Mymensingh, Bangladesh during November 2022 to July 2023. The studied samples (**Table S1**) were collected from five different locations including BAU residential area (BAURA), BAU Healthcare Center (BAUHCC), BAU Veterinary Teaching Hospital (BAUVTH), Poultry Market at BAU Sheshmor (PM), and Mymensingh Medical College Hospital (MMCH). A total of 110 samples (46 sources) from the study locations including BAURA = 15, BAUHCC = 20, BAUVTH = 22, PM = 30 and MMCH = 23 were collected (**Table S1**). Therefore, these samples belonged to three major categories including non-hospital environmental samples (BAURA; 15 samples), hospital-based clinical samples (BAUHCC, BAUVTH and MMHC; 65 samples) and poultry samples (PM, 30 samples). These samples comprising drain water, sewage, soil, and samples from hospital environments and poultry markets (**Table S1**), were individually placed in sterile test tubes with 3 – 4 mL nutrient broth, labelled and transported to the laboratory. A single loopful of overnight culture grown in nutrient broth was streaked over cetrimide agar (HiMedia, India) and aerobically incubated overnight at 37°C. Colonies with distinguishing characteristics, such as greenish color, were assumed to be *Pseudomonas* genus (n = 89 isolates). The genus-specific identification of *P. aeruginosa* typically involved a series of standard bacteriological methods including gram staining and an array of biochemical assays, including sugar fermentation, Voges-Proskauer, indole, and catalase tests [4, 17].

### 2.2. Molecular identification of *P. aeruginosa*

*Pseudomonas* isolates (n = 89) were then molecularly identified using genus- and species-specific polymerase chain reaction (PCR) assays. Genomic DNA from overnight culture by boiled DNA extraction method using commercial DNA extraction kit, QIAamp DNA Mini Kit (QIAGEN, Hilden, Germany). Quality and quantity of the extracted DNA were measured using a NanoDrop ND-2000 spectrophotometer (Thermo Fisher Scientific, Waltham, MA). DNA extracts with A260/280 and A260/230 ratios of ∼ 1.80 and 2.00 to 2.20, respectively, were considered as high-purity DNA samples [18] and stored at -20°C prior to PCR amplification [19, 20]. To amplify the 16S rRNA sequences of *Pseudomonas*, a set previously designed primer set was used (**Table 1**). PCR amplification of *16s_ Pseudo* for the detection of *Pseudomonas* genus [21] and *Pseudo_aeru* for the detection of *P. aeruginosa* species [22, 23] was performed on all phenotypically identified isolates of *P. aeruginosa*. The PCR condition against these primer sets for the amplification of genus- and species-specific primers is shown in **Table S2**. Amplification of targeted DNA was carried out in a 20 µL reaction mixture, including 3 µL nuclease-free water, 10 µL 2X master mixture (Promega, Madison, WI, USA), 1 µL forward and reverse primers (for each) and 5 µL DNA template. PCR-positive controls consisted of *P. aeruginosa* genomic DNA previously confirmed for the target genes [21–23]. PCR-negative controls utilized non-template controls with PBS instead of genomic DNA. Finally, amplified PCR products underwent electrophoresis on a 1.5% agarose gel and visualized using an ultraviolet transilluminator (Biometra, Gottingen, Germany). A 100 bp DNA ladder (Promega, Madison, WI, USA) was used to validate the expected sizes of the amplified PCR products [24]. Finally, 22 isolates were confirmed as *P. aeruginosa* through species-specific PCR.

**Table 1.**
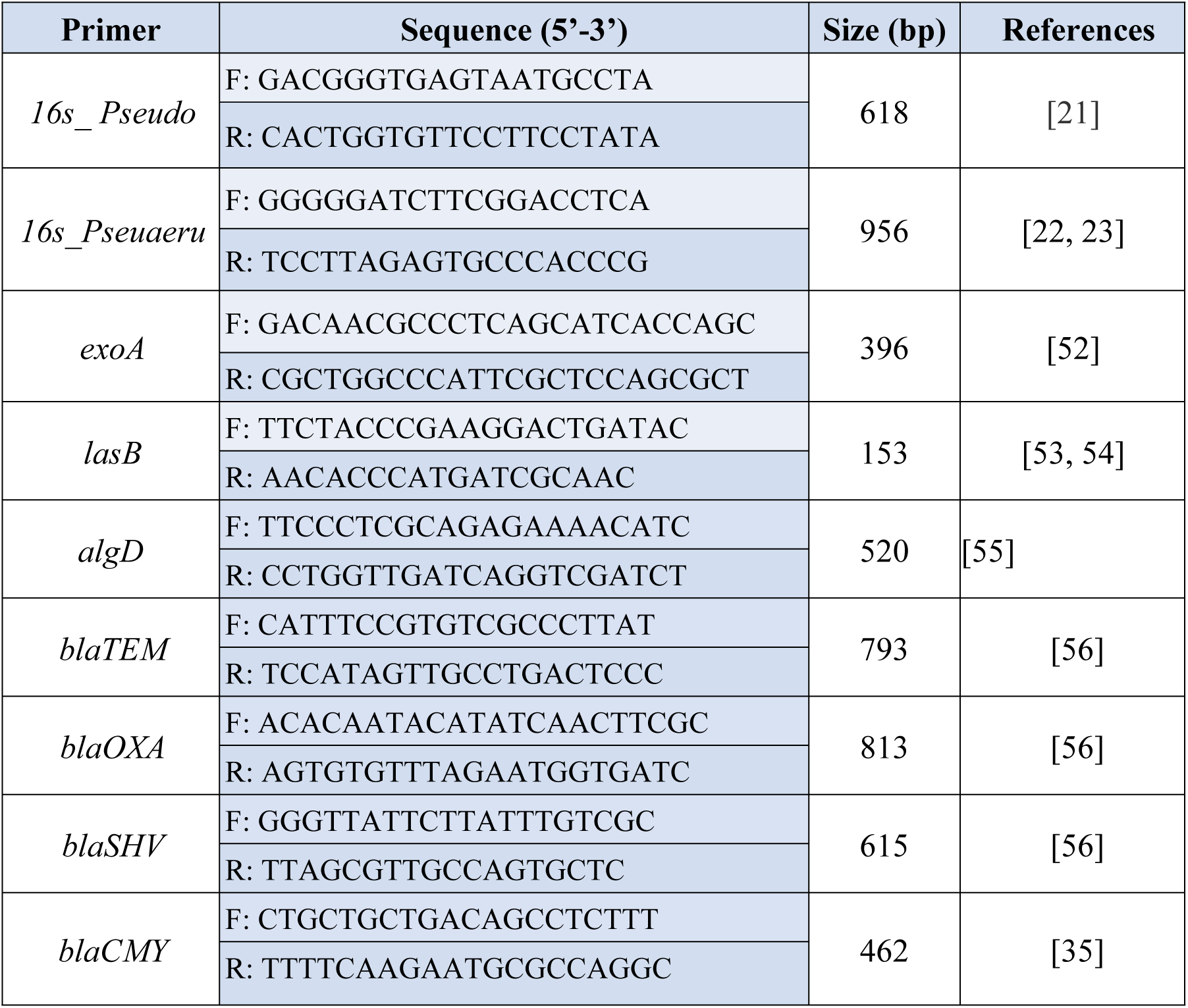
Primer sequences used in this study.

### 2.3. Antimicrobial susceptibility

The Kirby-Bauer agar disk diffusion method was used to test the susceptibility of the confirmed *P. aeruginosa* isolates to different antibiotics in accordance with the guidelines recommended by the Clinical and Laboratory Standards Institute (CLSI, 2021) to determine the sensitivity or susceptibility [25]. The antibiotic disks (Liofilchem^®^, Italy) used in this study included Ciprofloxacin (CIP, 5 g/disc), Gentamicin (GEN, 10 g/disc), Levofloxacin (LEV, 5 g/disc), Penicillin-G (P, 10 g/disc), Aztreonam (ATM, 30 g/disc), Imipenem (IMP, 10 g/disc), Amikacin (AK, 30 g/disc), Ceftazidime (CAZ, 30 g/disc), Doripenem (DOR, 10 g/disc), and Cefepime (CPM, 0 g/disc). Multidrug-resistant (MDR) isolates were those that were resistant to at least three antibiotic classes [26, 27]. The Multiple Antibiotic Resistance (MAR) index was derived by dividing the number of antibiotics an isolate resisted by the total number of antibiotics utilized [28].

### 2.4. Molecular detection of antibiotic resistant and virulence genes in *P. aeruginosa*

For the detection of the antibiotic resistant and virulence genes in the *P. aeruginosa* isolates, simplex PCR assays for beta-lactam antibiotics resistant genes (e.g., *blaTEM*, *blaCMY*, *blaSHV*, *blaOXA*), and virulent genes including *exoA, exoS, lasB,* and *algD* were performed with specific primers (**Table 1**). The PCR protocols utilized for detecting these genes were consistent with those described earlier in Section 2.2.

### 2.5. Statistical analysis

Data were entered into Microsoft Excel 2020^®^ (Microsoft Corp., Redmond, WA, USA) and analyzed using Excel and SPSS version 20 (IBM Corp., Armonk, NY, USA). The Pearson’s chi-square test was performed to compare the prevalence of *P. aeruginosa* in three different sample categories (e.g., clinical, environment and poultry). A prevalence percentage was calculated by dividing the number of positive samples for the given category by the total samples tested within that category [29, 30]. The prevalence formula was applied for determining prevalence percentage of *Pseudomonas* and *P. aeruginosa*. The AMR patterns, resistance, intermediate and sensitivity, and MAR index were calculated using the CLSI (2021) guideline using the cut-off as provided in the brochure of the manufacturer (Liofilchem^®^, Italy). Correlation coefficients for any of the two resistant antibiotics, association between phenotypic and genotypic resistance patterns, and phenotypic/genotypic resistance patterns and virulence genes of isolated *P. aeruginosa* was calculated using Pearson correlation tests. For the test, p < 0.05 was considered statistically significant.

## 3. Results

### 3.1 Prevalence of *P. aeruginosa*

From a total of 110 samples collected, 89 isolates were suspected to be *Pseudomonas* genus (80.90%) through culture and biochemical tests. However, the species-specific PCR confirmed 20% (22/110; 95% CI, 13.59-28.43) prevalence of *P. aeruginosa* in the studied samples. Non-hospital environment (from BAURA), hospital-based clinical (from BAUHCC, BAUVTH and MMCH) and poultry (PM) samples were found to harbor higher number of *P. aeruginosa*. The prevalence of *P. aeruginosa* was found to be the highest in the PM (30%, 95% CI, 12.6-47.4) followed by BAURA (26.67%, 95% CI, 12.4-27.6) and MMCH (13.85%, 95% CI, 0.5-22.5). Bivariate analysis exhibited that the hospital-based clinical (BAUHCC, BAUVTH and MMCH) samples had a significant correlation with both non-hospital environment samples and poultry market (PM) samples (Pearson correlation coefficient, ρ = 1; *p* <0.001) (**Table S3**).

### 3.2 Antibiogram profile of *P. aeruginosa*

The overall antibiogram profiles of isolated *P. aeruginosa* are presented in **Fig. 1**. All of the 22 *P. aeruginosa* isolates exhibited a concerning trend of antibiotic resistance. Notably, all isolates displayed 100% (95% CI, 85.13 – 100) resistance to Doripenem, Penicillin, Ceftazidime, Cefepime, and Imipenem, whereas resistance to Aztreonam was observed in 95.45% (95% CI, 78.20 – 99.76) of the isolates (**Fig. 1**). However, variable sensitivity was found against Gentamicin (77.27%, 95% CI, 56.56 – 89.87), and Levofloxacin (54.55%, 95% CI, 34.66 – 73.08), Amikacin (81.82%,95% CI, 61.48-92.69), and Ciprofloxacin (40.91%,95% CI, 23.25 – 61.26) (**Fig. 1**). The bivariate analysis demonstrated highly significant correlation between the antibiotic resistant isolates. A high positive significant correlation existed between resistance patterns of Levofloxacin and Ciprofloxacin (ρ= 0.641; *p*= 0.001), Amikacin and Gentamicin (ρ= 0.615; *p*= 0.002) and Levofloxacin and Gentamicin (ρ= 0.569; *p*= 0.006) (**Table 2**). Overall, MDR rate of the *P. aeruginosa* isolates was 100%. Among tested antibiotics, seven antibiotic resistance patterns were observed among the isolated *P. aeruginosa* isolates. Resistance pattern CAZ, CPM, IMP, DOR, ATM, P (Pattern No. 1) were the most observed (63.64%) among *P. aeruginosa* isolates. The MAR indices vary between 0.5 to 0.9 (**Table 3**).

**Fig. 1.**
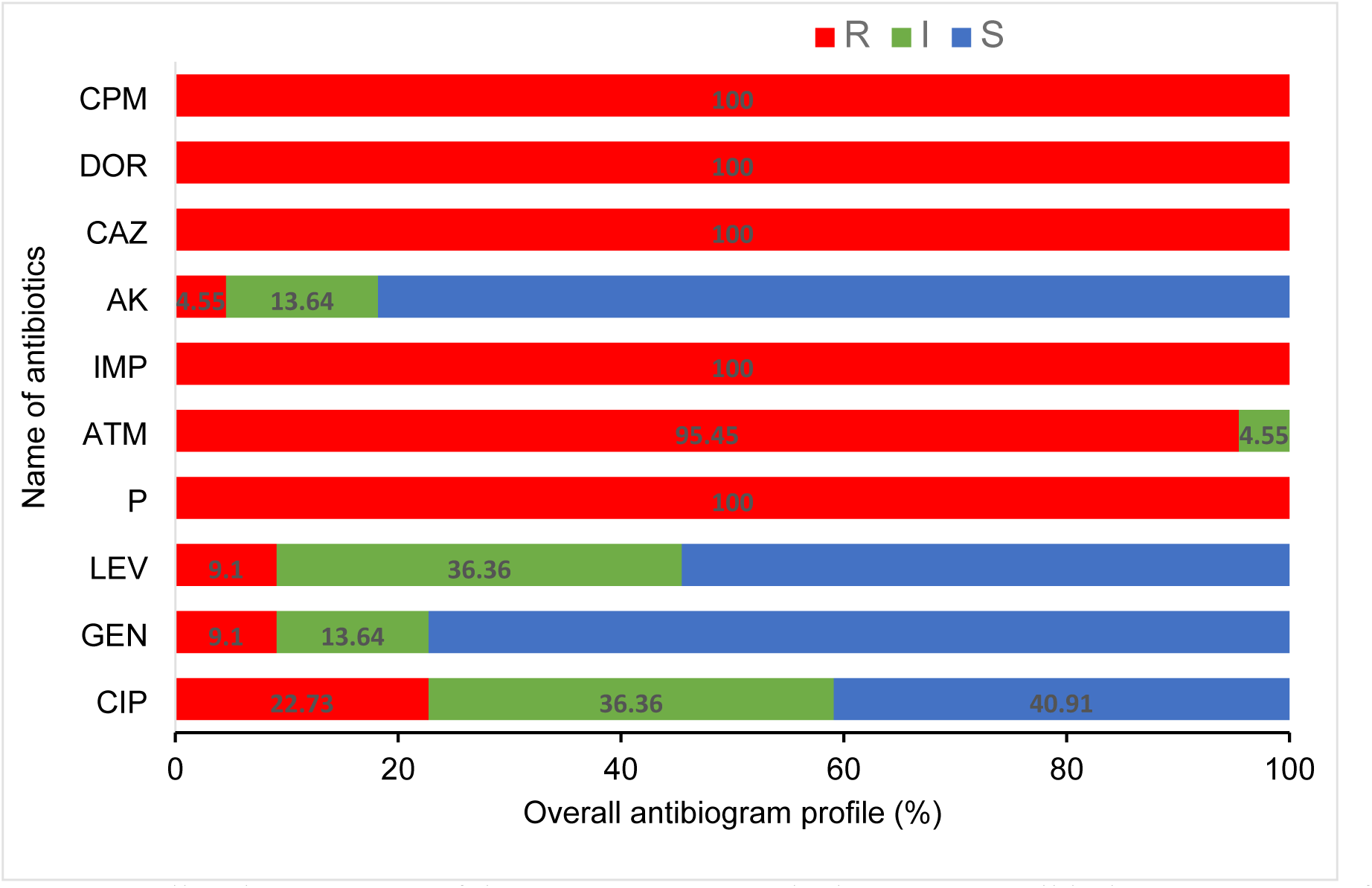
Overall resistance rates of the 22 *P. aeruginosa* isolates to 10 antibiotics. Percentage of R (Resistant, red), I (Intermediate resistant), and S (Susceptible) profiles are indicated for each antibiotic inside the bar chart. CIP: Ciprofloxacin, GEN: Gentamicin, LEV: Levofloxacin, P: Penicillin G, ATM: Aztreonam, IMP: Imipenem, AK: Amikacin, CAZ: Ceftazidime, DOR: Doripenem and CPM: Cefepime.

**Table 2.** Comparison of Pearson correlation coefficients between any of two antibiotics resistant to *P. aeruginosa*. **Legends**, A *p*-value less than 0.05 (p<0.05) was considered as significant; **, Correlation is significant at the 0.01 level (2-tailed); *, Correlation is significant at the 0.05 level (2-tailed). CIP: Ciprofloxacin, GEN: Gentamicin, LEV: Levofloxacin, ATM: Aztreonam, and AK: Amikacin.

|  |  | CIP | GEN | LEV | ATM | AK |
| --- | --- | --- | --- | --- | --- | --- |
| CIP | Pearson Correlation | 1 |  |  |  |  |
|  | Sig. (2-tailed) |  |  |  |  |  |
| GEN | Pearson Correlation | .396 | 1 |  |  |  |
|  | Sig. (2-tailed) | .068 |  |  |  |  |
| LEV | Pearson Correlation | .641** | .569** | 1 |  |  |
|  | Sig. (2-tailed) | .001 | .006 |  |  |  |
| ATM | Pearson Correlation | .230 | .110 | .182 | 1 |  |
|  | Sig. (2-tailed) | .303 | .626 | .419 |  |  |
| AK | Pearson Correlation | .330 | .615** | .171 | .096 | 1 |
|  | Sig. (2-tailed) | .134 | .002 | .447 | .671 |  |

**Table 3.** Multidrug-resistance patterns of *P. aeruginosa* isolates.

| Pattern No. | Antibiotic Resistance Patterns | No. of Antibiotics (Classes) | No. of MDR Isolates (%) | Overall No. of MDR Isolates (%) | MAR indices |
| --- | --- | --- | --- | --- | --- |
| 1 | CAZ, CPM, IMP, DOR, ATM, P | 6 (4) | 14 (63.64) | 22 (100) | 0.6 |
| 2 | CAZ, CPM, IMP, DOR, ATM, P, LEV | 7 (5) | 1 (4.55) |  | 0.7 |
| 3 | CIP, CAZ, CPM, IMP, DOR, ATM, P, LEV | 8 (5) | 1 (4.55) |  | 0.8 |
| 4 | CIP, CAZ, CPM, IMP, DOR, ATM, P | 7 (5) | 3 (13.64) |  | 0.7 |
| 5 | CIP, CAZ, GEN, CPM, IMP, DOR, ATM, P, LEV | 9 (6) | 1 (4.54) |  | 0.9 |
| 6 | CAZ, CPM, IMP, DOR, P | 5 (3) | 1 (4.54) |  | 0.5 |
| 7 | CAZ, GEN, CPM, IMP, DOR, ATM, AK, P | 8 (5) | 1 (4.54) |  | 0.8 |

### 3.3. Antibiotic resistance genes of *P. aeruginosa*

Of the tested isolates (n = 22), 19 (86.36%, 95% CI, 66.66 – 95.25); 6 (27.27%, 95% CI, 13.15 – 48.15); 4 (18.18%, 95% CI, 7.30 – 38.51); and 1 (4.55%, 95% CI, 0.23 – 21.79) were positive for *blaTEM*, *blaCMY*, *blaSHV*, and *blaOXA*, respectively, which are responsible for beta-lactam antibiotics resistance in *P. aeruginosa*. Among the resistance genes harboring isolates, 9.09% (95% CI, 1.61-27.81) isolates were positive for the *blaTEM*, *blaSHV*, and *blaOXA* genes; 4.54% (95% CI, 0.23-21.79) were for *blaTEM*, *blaCMY*, and *blaOXA* genes; 9.09% (95% CI, 1.61– 27.81) for the *blaTEM*, *blaCMY*, *blaSHV*, and *blaOXA* genes; 13.63% (95% CI, 4.74 – 33.33) for the *blaTEM* and *blaCMY* genes; and 50% (95% CI, 30.72 – 69.27) were found to carry a single resistance gene, *blaTEM* (**Fig. 2**). By comparing the phenotypic and genotypic resistance pattern of the *P. aeruginosa* isolates, we found that the Levofloxacin and Gentamycin resistant isolates had significantly higher amount of *blaCMY* and *blaOXA* than the other two checked genes (*blaTEM* and *blaSHV*) (**Table 4**).

**Fig. 2.**
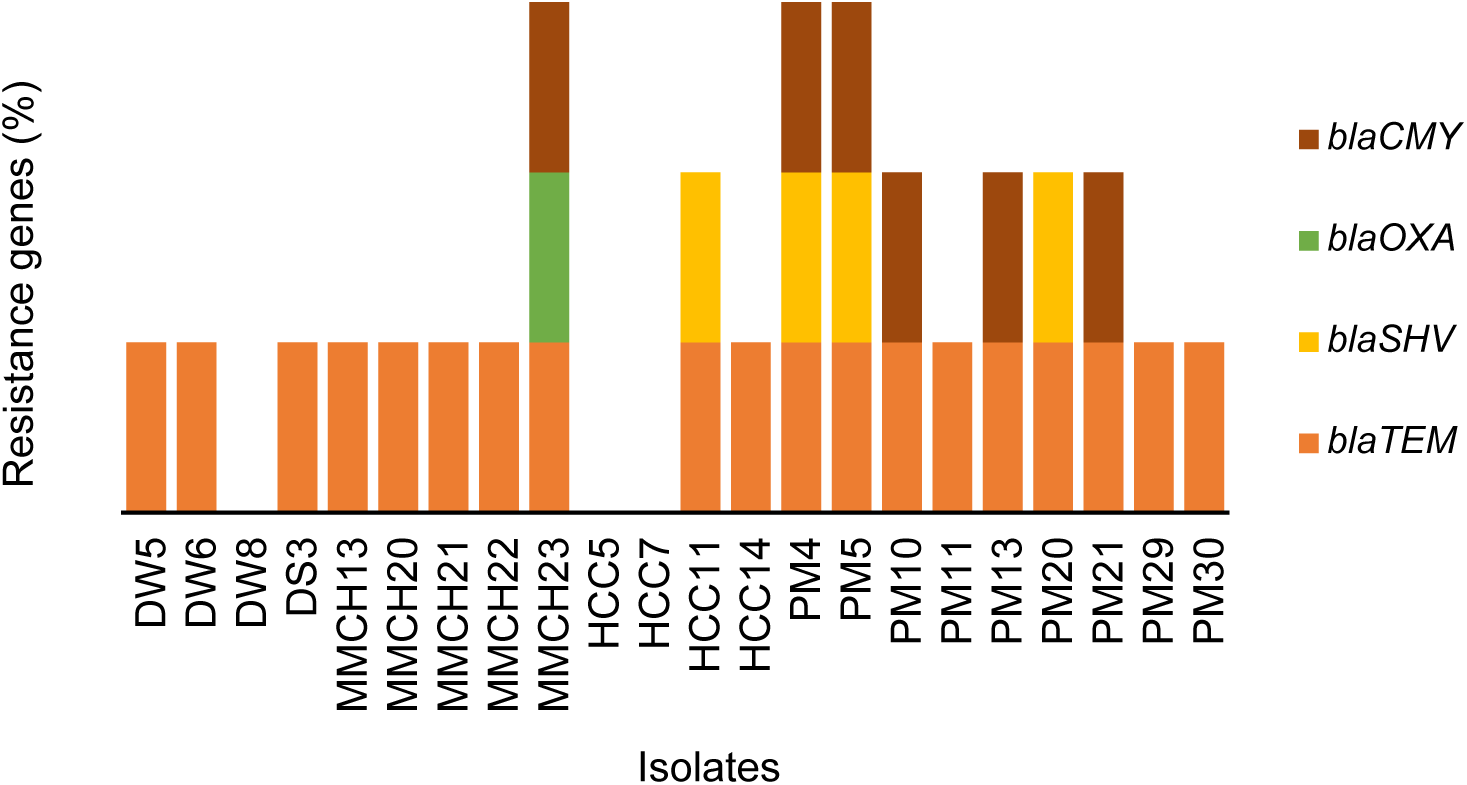
Distribution of antimicrobial resistance (β-lactamase) genes (ARGs) in *P. aeruginosa* currently circulating in clinical, environmental and poultry sources in Bangladesh.

**Table 4.** Association between phenotypic and genotypic resistance patterns of isolated *P. aeruginosa*. Each subscript letter denotes a subset of genes categories whose column proportions do not differ significantly from each other at the 0.05 level. NA: Not applicable.

| Resistance | Genotypic |  |  |  |  |  |
| --- | --- | --- | --- | --- | --- | --- |
|  | Antibiotics | No (%) of<br>*bla*TEM<br>(n=19) | No (%) of<br>*bla*CMY<br>(n=6) | No (%) of<br>*bla*SHV<br>(n=4) | No (%) of<br>*bla*OXA<br>(n=1) | *p*-value |
| Phenotypic | CPM | 19 (100<sup>a</sup>) | 6 (100<sup>a</sup>) | 4 (100<sup>a</sup>) | 1 (100<sup>a</sup>) | NA |
|  | DOR | 19 (100<sup>a</sup>) | 6 (100<sup>a</sup>) | 4 (100<sup>a</sup>) | 1 (100<sup>a</sup>) | NA |
|  | CAZ | 19 (100<sup>a</sup>) | 6 (100<sup>a</sup>) | 4 (100<sup>a</sup>) | 1 (100<sup>a</sup>) | NA |
|  | AK | 1(5.3<sup>a</sup>) | 0(0<sup>a</sup>) | 0(0<sup>a</sup>) | 0(0<sup>a</sup>) | 0.897 |
|  | IMP | 19(100<sup>a</sup>) | 6(100<sup>a</sup>) | 4(100<sup>a</sup>) | 1(100<sup>a</sup>) | NA |
|  | ATM | 18(94.7<sup>a</sup>) | 6 | 3(75<sup>a</sup>) | 1 | 0.437 |
|  | P | 19(100<sup>a</sup>) | 6(100<sup>a</sup>) | 4(100<sup>a</sup>) | 1(100<sup>a</sup>) | NA |
|  | LEV | 1(5.3<sup>a</sup>) | 1(16.7<sup>b</sup>) | 0(0<sup>a</sup>) | 1(100<sup>b</sup>) | 0.017 |
|  | GEN | 2(10.5<sup>a</sup>) | 1(16.7<sup>b</sup>) | 0(0<sup>a</sup>) | 1(100<sup>b</sup>) | 0.050 |
|  | CIP | 4(21.1<sup>a</sup>) | 2(33.3<sup>a</sup>) | 1(25<sup>a</sup>) | 1(100<sup>a</sup>) | 0.362 |

### 3.4. Virulence genes in *P. aeruginosa*

Gene-specific simplex PCR was used to screen the virulence genes in 22 *P. aeruginosa* isolates. Among these isolates, 31.81% (7/22, 95% CI, 16.36-52.68) were positive for the *algD* gene and 9.1% (2/22, 95% CI, 1.61-27.81) were positive for *lasB* gene. However, *exoA* gene was detected only in 4.55% (1/22, 95% CI, 0.23-21.79) *P. aeruginosa* isolates (**Fig. 3**), while *exoS* gene was absent in the study isolates. Moreover, 9.1% (2/22, 95% CI, 1.61-27.81) isolates were found to carry both *lasB* and *algD* genes, whereas 4.54% (1/22, 95% CI, 0.23-21.79) isolates harbored both *algD* and *exoA* genes. Additionally, by analyzing the association between resistance patterns (phenotypic or genotypic) and virulence profile (genotypic) of the *P. aeruginosa* isolates, we found no significant association (**Tables S4 and S5**).

**Fig. 3.**
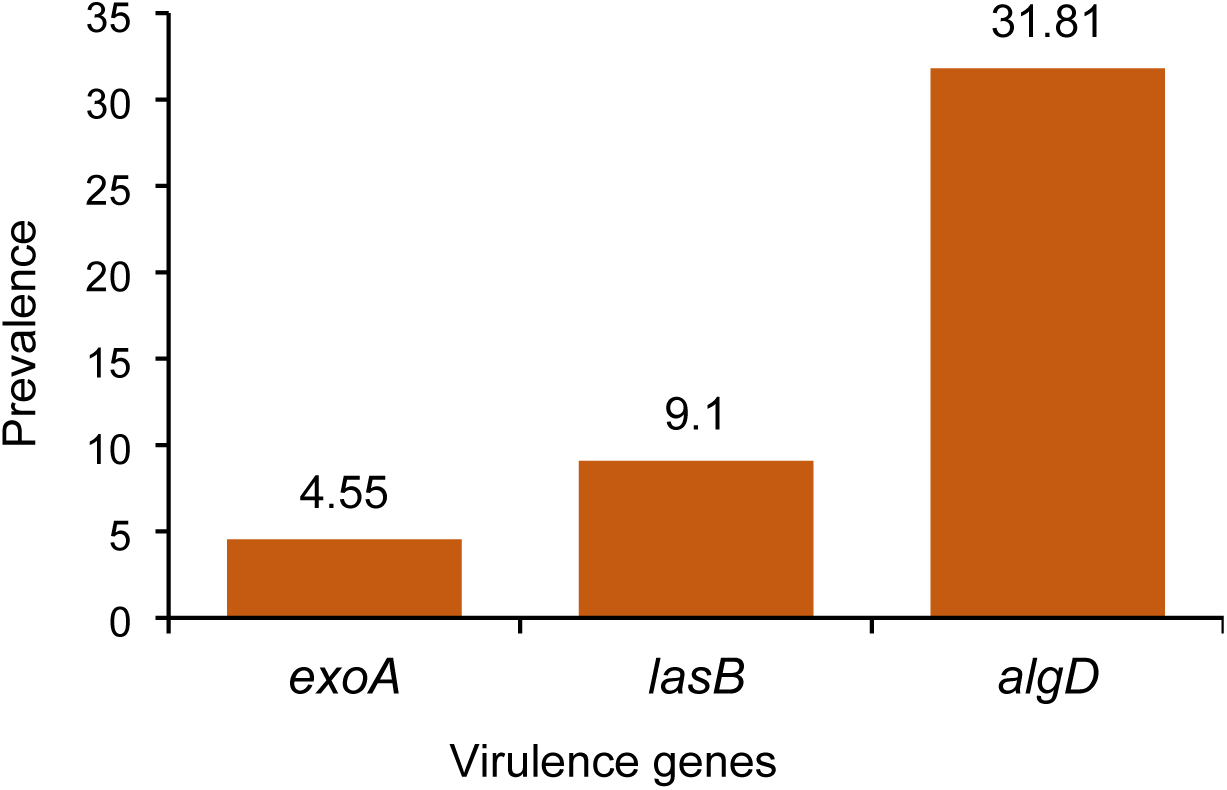
Prevalence of virulence genes in *P. aeruginosa* currently circulating in clinical, environmental and poultry sources in Bangladesh.

## 4. Discussion

*Pseudomonas aeruginosa* is considered a significant cause of nosocomial or healthcare-associated infections (HAI) and is classified as a priority critical pathogen of global concern by various health organizations [31]. As an opportunistic bacterial pathogen, *P. aeruginosa* has been linked with a range of human and animal infections [32]. *P. aeruginosa* is ubiquitous in the environment (soil and water) and capable to produce disease in individuals with weakened or compromised immune systems [33]. Despite breakthroughs in medical and surgical treatment, as well as the introduction of a wide range of antimicrobial drugs in livestock and agricultural, the number of resistant isolates in outdoor surface water has increased considerably [34]. Reports from different countries are available describing the detection of *P. aeruginosa* from various environmental sources [4, 5, 33, 35]. However, sufficient information is unavailable on the environmental distribution of potential pathogenic MDR *P. aeruginosa* in Bangladesh. Previous studies in Bangladesh have focused on clinical isolates, healthcare workers’ mobile phones, and even the genomic diversity of MDR *P. aeruginosa* from burn patients [4, 36, 37]. Therefore, this study was designed to determine the antibiotic resistance profiles and virulence genes distribution in *P. aeruginosa* circulating in different environments of Bangladesh. We analyzed 110 samples belonged to three major categories e.g., non-hospital environment (BAURA), hospital-based clinical (BAUHCC, BAUVTH and MMHC) and poultry market (PM) from five distinct locations of Mymensingh division of Bangladesh. These diverse study areas collectively provided a comprehensive overview of the prevalence and distribution of MDR *P. aeruginosa* and its virulence profiles within distinct environmental contexts, contributing to a holistic understanding of the potential public health significance of these bacteria in the selected region. The overall prevalence of *P. aeruginosa* was 20.0%. However, based on locations and sample categories, 26.67%, 13.85%, and 30.0% were reported in non-hospital environment samples (drain water and drain sewages) of BAURA, clinical samples (BAUHCC, BAUVTH and MMCH) and poultry (PM) samples, respectively. The highest isolation rates of *P. aeruginosa* (30.0%) was found in the PM samples. In Bangladesh, the average isolation rate for *P. aeruginosa* from diverse surface waters was reported to be 61.5%, which is greater than in its neighboring nation, India (45.45%) [38]. The isolation rate of various water samples may fluctuate depending on the source and size of the sample, as well as the time and length of sample collection [3, 35, 39]. Previously, Algammal et al. reported 28.3% prevalence of *P. aeruginosa* in broiler chickens in Egypt [40] whereas El-Tawab et al. found that 34% of the tested frozen fish possessed *P. aeruginosa* [41]. Detection of *P. aeruginosa* in poultry and associated environment is alarming because of the possibilities of transmission of these pathogens to human.

In this study, the most commonly used antibacterial drugs in Bangladesh were tested against *P. aeruginosa* isolates to determine their resistance pattern. We found that 100% *P. aeruginosa* isolates were resistant to multiple antibiotics such as Penicillin, Ceftazidime, Cefepime, Doripenem and Imipenem. In addition, 95.45% isolates were resistant to Aztreonam. However, the study isolates showed high to moderate level (44.0 to 78.0%) sensitivity against Amikacin, Gentamicin and Ciprofloxacin. These results corroborated with several earlier studies that investigated antimicrobial susceptibility profile in P. aeruginosa isolated from clinical samples [4], environmental samples [37] and poultry samples [42]. Consistent findings across various studies examining the antimicrobial susceptibility profile of *P. aeruginosa* isolated from diverse sources (clinical, environmental, and animal samples) can lend support to the understanding of the AMR patterns and prevalence of MDR strains within this bacterium. Such cumulative evidence aids in comprehending the overall resistance landscape and guides the development of effective strategies for managing infections caused by *P. aeruginosa*. We found that 100.0% isolates of the *P. aeruginosa* were MDR isolates (resistant to > 5 antibiotics). Furthermore, the multiple antibiotic resistance (MAR) indices for the *P. aeruginosa* isolates ranged from 0.5 to 0.9 indicating an increased likelihood of these isolates being exposed to various antibiotics or antimicrobial agents, potentially contributing to the development of MDR, posing significant concerns for public health due to the challenges in treating infections caused by these organisms. Indeed, a MAR index greater than 0.2 is indicative of potential contamination from sources where antibiotics are frequently utilized [43].

We also investigated the occurrence of several ARGs which could aid in designing effective treatment strategies and surveillance programs to control the spread of antibiotic-resistant strains of *P. aeruginosa*. In this study, *blaTEM* gene was found to be predominating in 86.36% isolates. This high frequency of *blaTEM* is however, consistent with the findings of Hosu et al. who reported a 79.3% prevalence of *blaTEM* in clinical *P. aeruginosa* isolates, indicating its widespread dispersion in healthcare settings [11, 44]. the second most prevalent ARG was *blaCMY* which was detected in about 27.0% isolates. Recently, Ejikeugwu et al. reported that around 42.0% *P. aeruginosa* isolates from clinical samples harbored *blaCMY* gene [45], which is a bit higher than our findings. This observation is incongruent with the findings of Mohamed et al. who detected a far higher incidence of *blaCMY* in *P. aeruginosa* isolates derived from chickens, indicating possible sources of resistance in non-clinical settings [46]. Although, *blaSHV*, and *blaOXA* were identified as the less abundant ARG in *P. aeruginosa* isolates (18.0% and 4.55%, respectively), these ARGs were found in different combination. The detection of *blaSHV*, and *blaOXA* genes in *P. aeruginosa* isolates aligns with findings from previous studies [4, 47, 48], further highlighting the significance of these genes in healthcare settings and their association with nosocomial infections. Their presence underscores the potential for these bacteria to cause severe infections and emphasizes the need for stringent infection control measures and judicious use of antibiotics to mitigate the emergence and spread of multidrug-resistant strains The difference between the results of this study and other investigations might be attributed to differences in various circumstances comprising incubators or to commonly occurring hyper-mutation among *P. aeruginosa* strains exhibiting varied antibiotic resistance. Furthermore, antibiotic-resistant bacteria can swiftly spread across the food chain and cause the majority of public health problems [34, 36].

One of the hallmark findings of this study is the identification several virulence genes in the *P. aeruginosa* isolates that might contribute to the pathogenicity of this bacterium in multiple hosts. The presence of the *algD* and *lasB* genes in around 32.0% and 9.0% of the *P. aeruginosa* isolates, respectively, suggests the potential virulence of these strains. The *algD* gene is associated with alginate production, which contributes to the formation of biofilms and enhances the bacteria’s resistance to host immune responses and antibiotics [49, 50]. The *lasB* gene encodes for elastase, an enzyme involved in tissue damage and immune evasion [51]. On the other hand, the low prevalence of the *exoA* gene (4.55%) and the absence of the *exoS* gene indicate the variability in the distribution of virulence factors among these isolates. The *exoA* gene encodes exotoxin A, a cytotoxin known to interfere with host cell function, contributing to cytotoxicity and the establishment of infections [52]. Therefore, the differential presence and absence of these virulence genes among the isolates underscore the heterogeneity of *P. aeruginosa* strains and their potential variation in virulence and pathogenicity.

## 5. Conclusions

To the best of our knowledge this is the first report on detection of MDR *P. aeruginosa* from various environmental samples including non-hospital residential environment (BAURA), hospital-based clinical (human and animal hospitals) and poultry markets samples for the first time in Bangladesh. We found that *P. aeruginosa* is prevalent in these diverse group of samples at 20.0%, and highest prevalence was recorded in poultry samples (30.0%). Despite having discrepancy in prevalence according to sample types, 100.0% *P. aeruginosa* isolates showed resistance against at least five antibiotics tested. Additionally, identified MDR *P. aeruginosa* were found to harbor beta-lactamase genes such as *blaTEM*, *blaCMY*, *blaSHV*, and *blaOXA* and virulence genes including *exoA, lasB*, and *algD*. Occurrence of MDR in *P. aeruginosa* is very alarming, since human and other animals can easily pickup these potential pathogens from the environment to get infection. Routine surveillance and practice of regular disinfection of environmental surfaces could be adopted rigorously to reduce their load in the environments. Indeed, there are several promising avenues for further research on the *P. aeruginosa* isolates obtained in this study. Conducting further investigations using advanced techniques such as whole-genome-based phylogenetic analysis, epidemiological source tracking, sequence typing of isolated *P. aeruginosa* strains, and determining the minimum inhibitory concentration (MIC) of antibiotics, resistome, and pathogenicity profiles would significantly contribute to a comprehensive understanding of various aspects related to *P. aeruginosa* infections. Leveraging a larger cohort of samples in these investigations would increase the robustness and representativeness of the findings, leading to more reliable conclusions and facilitating the development of more effective strategies for infection control, treatment, and management of *P. aeruginosa* infections.

## Author’s Contributions

Raihana Islam: Sample collection, Raihana Islam, Farhana Binte Ferdous, Nowshad Atique Asif, Md. Liton Rana: Experiment, Data curation, Formal analysis, Methodology, Writing – original draft, M. Nazmul Hoque: Methodology, Validation, Visualization, review and editing, Mahbubul Pratik Siddique: Project administration, Supervision, Writing – review and editing, and Md. Tanvir Rahman: Conceptualization, Funding acquisition, Project administration, Supervision, Writing – review and editing.

## Acknowledgments

The authors would like to thank the authority who provided us with the samples from diverse environment to the support the research.

## Consent for publication

Not applicable.

